# scSAGA: Single-cell Sampled Gromov Wasserstein Alignment for Scalable and Memory-efficient Integration of Multi-modal Single Cell Data

**DOI:** 10.64898/2026.03.26.714573

**Authors:** Swethasree Bhattaram, Sriram Chockalingam, Maneesha Aluru, Srinivas Aluru

## Abstract

**Motivation:** Several different methods exist for multimodal integration of single cell RNA-seq (scRNA-seq) and chromatin accessibility (scATAC-seq) data. However, these methods either suffer from quadratic memory and runtime complexity, which hinders their applicability to large datasets, or trade off geometric fidelity for efficiency, which limits performance when modalities have disjoint features. Consequently, there is no existing framework that simultaneously preserves manifold structure and scales to organism-wide multimodal single cell datasets.

**Results:** We present scSAGA (Single-Cell Sampled Gromov–Wasserstein Alignment), a geometry-preserving, scalable and memory-efficient method designed for integration of paired and unpaired scRNA-seq and scATAC-seq datasets. scSAGA combines (i) sparse *k*NN graph geometry with on-demand geodesic distances, (ii) plan-guided sampled Gromov– Wasserstein optimization, and (iii) a matrix-free joint embedding computed with sparse iterative linear algebra. Across paired and unpaired benchmark datasets from various organisms including Human PBMC and BMMC, mouse Alzheimer’s brain, Zebrafish, and *Arabidopsis* root, scSAGA achieves significantly improved one-to-one matching accuracy and/or modality mixing relative to well-established methods such as Pamona, SCOT, Seurat, and LIGER, while also scaling to integrations exceeding one million cells with near-linear growth in runtime and memory. Furthermore, scSAGA yields stronger downstream clustering of the integrated multimodal data, resulting in more coherent clusters for cell-type identification. scSAGA is thus the first geometry-preserving, memory-efficient optimal transport framework capable of accurate and scalable single-cell multimodal integration.

**Code and Data Availability:** Code is available at https://github.com/AluruLab/scSAGA. The full list of datasets is listed in the supplementary table 1.

## Introduction

High-throughput single cell multimodal assays profile complementary molecular layers from the same tissue - most commonly transcriptome (scRNA-seq) and chromatin accessibility (scATAC-seq), enabling cell-resolved views of development, disease, and perturbation [6]. A core computational challenge in the integrative analysis of such datasets is *multimodal integration* while preserving intra-modality structure, i.e., aligning cells across modalities when feature spaces are unmatched (genes vs. peaks) and measurements may be unpaired [22]. In practice, integration must also handle partial population overlap, batch/study effects, and modern dataset sizes that routinely exceed 100, 000 to a million cells.

**Table 1.** Integration accuracy (1:1) on the paired Human PBMC Granulocytes dataset for different integration sizes.

| # Cells | scSAGA | Pamona | SCOTv2 | LIGER | Seurat v5 |
| --- | --- | --- | --- | --- | --- |
| 600 | **0.9970** | 0.8829 | 0.2000 | 0.7812 | 0.9300 |
| 2000 | **0.9910** | 0.9138 | 0.0970 | 0.6513 | 0.8100 |
| 10000 | **0.9882** | 0.8135 | 0.0338 | 0.7512 | 0.8560 |
| 22602 | **0.9536** | 0.7810 | 0.0500 | 0.6514 | 0.7713 |

Currently available multimodal integration methods follow one of two broad strategies - shared-feature alignment and geometry-based alignment. *Shared-representation* approaches learn a joint latent space using linear or factor models (e.g., Seurat, Harmony, LIGER). However, in cross-modality settings they often rely on proxy shared features such as gene activity, which can introduce modeling choices that affect the resulting geometry [5, 21, 15, 25, 23]. *Geometry-based alignment* or the *optimal transport-based* approaches instead match modalities using intra-modality structure rather than raw features. Here, the Gromov–Wasserstein (GW) optimal transport method is attractive because it aligns datasets via their *intra-domain* distances through a probabilistic coupling, without requiring feature correspondences [18, 10, 7, 11, 20]. The GW method treats each dataset as a metric-measure space and seeks a coupling that best preserves intra-domain geodesic distances (e.g., along kNN graphs). For cross-modality integration, this avoids fragile pseudo shared features and admits principled generalizations to partial and unbalanced settings when populations overlap only in part, thus making this framework practical for multimodal integration of single cell data.

Despite these advantages, existing GW-based single-cell methods (SCOT, Pamona, SCOTv2) remain difficult to scale because of two systemic bottlenecks. First, they typically precompute and store dense all-pairs intra-domain geodesic distances, which requires 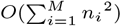memory, where *M* is the number of datasets in the integration and *n*_*i*_ is the size of the *i*^*th*^ dataset. Second, they assemble and optimize over the full GW cost, which is quadratic because of all pairwise comparisons across both datasets. Together, these choices become prohibitive in runtime/memory costs beyond a few thousand cells, and also impede simultaneous alignment of multiple datasets in one pass. [10, 11, 7].

To address these issues, we developed **scSAGA** (**s**ingle-**c**ell **sa**mpled **g**romov–wasserstein **a**lignment), a de novo geometry-based framework that retains GW alignment while removing its principal scalability bottlenecks. Our method (i) avoids dense distance matrices via on-demand geodesic queries on sparse *k*NN graphs, (ii) approximates GW costs using plan-guided sampling, and (iii) computes a multi-dataset joint embedding in a matrix-free manner from couplings to an anchor dataset. We show that scSAGA outperforms existing multimodal scRNA-seq and scATAC-seq integration methods including previous optimal transport based methods, and achieves strong accuracy and alignment quality across both paired and unpaired benchmark datasets, while scaling to million+ cells in a memory-efficient manner.

## Methods

GW optimal transport provides a rigorous framework to align multi-modal single cell datasets by relying on cross-domain structure rather than shared features. We formally define the GW optimal transport problem for single-cell datasets below.

### Background: GW Optimal Transport

Let *X*_1_ and *X*_2_ denote datasets of *n*_1_ and *n*_2_ cells, respectively, obtained from two different modalities (e.g., scRNA and scATAC). Let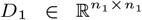 and 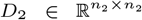 be the corresponding pairwise distance matrices. Entry 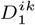 represents the entry in row *i* and column *k* in *D*_1_, and captures the respective cell–cell similarity within the respective modality, typically computed based on PCA or latent semantic indices.

GW-based alignment methods aim to learn a **transport plan** 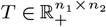, where the entry *T*^*ij*^ defines the probability mass from cell *i* in modality *X*_1_ is assigned, i.e., *transported*, to cell *j* in modality *X*_2_. A higher *T*^*ij*^ indicates that the two cells are likely counterparts across modalities. In order to identify a plan *T* that preserves local geometry of the modalities, GW distance is defined as the following optimization problem:

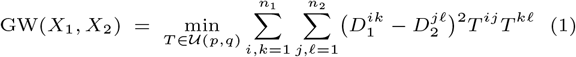

where 𝒰 (*p, q*) is the set of nonnegative matrices, whose row and column sums are the vectors *p* and *q* respectively, which sum to 1 (usually defining a uniform distribution). The GW objective ensures that two cells are matched if their relative distances to other cells are similar, even if features differ between modalities [18, 20, 24]. To enable efficient solutions to (1), standard GW methods assume that both the datasets have the same number of cells (*n*_1_ = *n*_2_). *Partial* GW overcomes this limitation by matching only a shared fraction of the total mass and allowing unmatched cells via a virtual (sink) mass with a large penalty [7, 11]. The resulting *T* is a soft correspondence matrix where confident cross-modality matches receive high mass.

Multi-modal integration with *M* datasets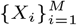of sizes *n*_1_, *n*_2_, …, *n*_*M*_, respectively, is typically reduced to a set of *M* − 1 pairwise partial-GW problems by selecting an *anchor* dataset *X*_*a*_ and computing 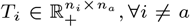. This approach avoids solving 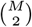 GW problems and mirrors anchor-hub strategies used in most multi-dataset integration pipelines [7, 11].

- **Distance-side bottleneck:** Computing and storing dense distance matrices i.e., *D*_*i*_, 1 ≤ *i* ≤ *M*, requires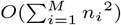memory for *M* datasets, where *n*_*i*_ is the size of the ith dataset, which is infeasible for tens to hundreds of thousands of cells.
- **Objective-side bottleneck:** When solving eq. 1, the dominant term in the gradient of the objective function with respect to *T* is *D*_1_*TD*_2_, computing which takes *O*(*n*_1_*n*_2_(*n*_1_+ *n*_2_)) time and consumes *O*(*n*_1_*n*_2_) memory.

In the balanced case, where all the *M* datasets have the same number of cells, say *n*, i.e., *n* = *n*_1_ = … = *n*_*M*_, solving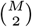GW problems costs *O*(*M* ^2^*n*^3^) compute time and *O*(*Mn*^2^) memory, making standard approaches impractical for organism-wide multimodal integration.

### scSAGA approach for scalable GW

Our method **scSAGA**, outlined in figure 1, addresses these challenges with the following novel strategies:

**Fig. 1.**
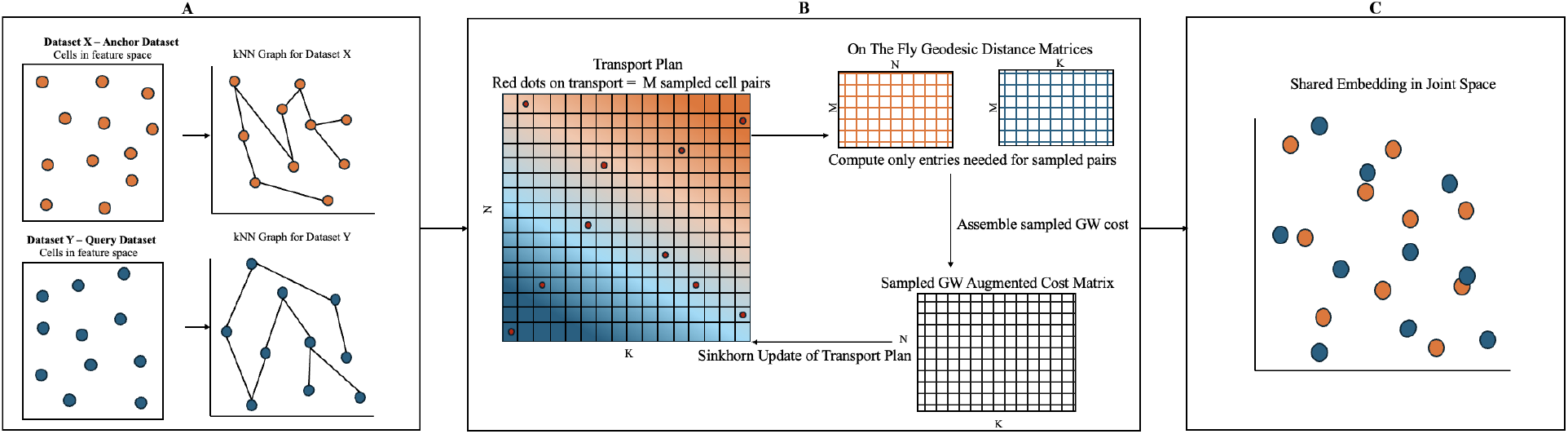
scSAGA workflow. (A) Sparse neighborhood construction. Cells are converted into a sparse *k*NN graph to capture intra-dataset local geometry. **(B) Sampled partial-GW alignment to an anchor**. A soft transport plan is iteratively refined: cell pairs (red dots) are sampled from the current plan, geodesic-distance entries are computed on-the-fly, a sampled augmented GW cost is assembled, and a Sinkhorn update produces a refined plan. **(C) Shared embedding**. Using the learned transport plans with graph-based regularization, cells from all datasets are mapped into a common low-dimensional space for direct cross-dataset comparison.

1. **Sparse geometry with on-the-fly geodesics**. Each dataset *X*_*i*_ is represented by a sparse *k*-nearest-neighbor (kNN) graph. Distances are computed as graph geodesics *only when needed*, avoiding dense 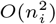distance storage (See Figure 1A).
2. **Plan-guided sampled GW**. In each iteration, the GW objective is approximated using a small set of informative index pairs sampled from the current plan, focusing computation where the plan has high mass [14] (See Figure 1B).
3. **Matrix-free joint embedding**. After estimating couplings to the anchor, the joint embedding is computed using iterative linear algebra on sparse operators (matrix–vector products and iterative solves), avoiding dense matrix factorization (See Figure 1C).

### *scSAGA* Algorithm

Algorithm 1 describes the **scSAGA** method for integrating *M* datasets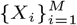. Without loss of generality, we assume that the first dataset *X*_1_ to be the anchor dataset. The algorithm proceeds in three phases: (i) build a sparse neighborhood graph for each dataset, (ii) estimate a partial-GW coupling from every non-anchor dataset to the anchor dataset, and (iii) compute a single joint embedding for all cells. Each dataset *X*_*i*_ is represented in a latent space of dimension *r*_*i*_, i.e., 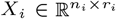Given the anchor dataset *X*_1_ with *n*_1_ cells, scSAGA outputs (a) GW plans 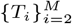with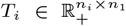, and (b) embeddings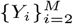 with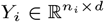, where *d* is the dimension of the common embedding space. A visual representation of the algorithm is shown in Figure 1.

### Sparse graphs and geodesic distances (lines 1–4)

For each dataset *i*, we build a *k*NN graph *G*_*i*_ in the latent space and ensure that the graph is connected by adding low cost inter-component edges when needed. *G*_*i*_ provides a memory-light representation of intra-dataset geometry (sparse weights 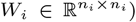, and enables shortest-path (geodesic) distance queries without fully computing and storing the dense *n*_*i*_ × *n*_*i*_ distance matrix in memory.

### Sampled partial-GW plans to the anchor (lines 5–13)

For a non-anchor dataset *D*_*i*_, 1 *< i* ≤ *M*, let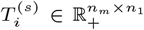 be the current transport plan at iteration *s*. At each iteration, we sample *B* index pairs 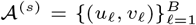from 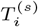, and compute only the required geodesic distance for the sampled columns on *G*_*i*_ and *G*_*a*_. We evaluate the (*b, c*) entry in the sampled GW cost matrix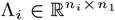as

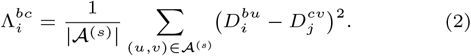

### Solution to Partial GW (lines 14– 23)

To allow unmatched cells, we augment Λ_*i*_ with a virtual row and column, yielding 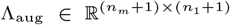with virtual-cell penalty *ζ*. We use partial marginals *p*^′^, *q*^′^ so that only a total mass *ρ* is required to match between real cells. We then solve the entropically regularized problem with Sinkhorn iterations [9, 12] (line 16), and remove the virtual cells to obtain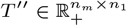. After the plan is updated with damping 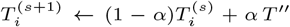(line 17), we proceed to the next iteration, unless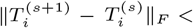 `tol`. Repeating the alignment for each *i* (1 *< i* ≤ *M*) yields couplings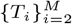.

### Matrix-free joint embedding (lines 24–32)

After all the transport plans from each dataset to the anchor are computed, we construct a single low-dimensional space where all cells can be compared. Intuitively, the embedding is built to satisfy two requirements: (i) cells that are neighbors within the same dataset (on the *k*NN graph) should remain close, and (ii) cells that receive high transport mass across datasets should be as close as possible. To enforce (i), we use the *k*NN graphs *G*_*i*_ (1 ≤ *i* ≤ *M*) to define a smoothness penalty (via the graph Laplacian). To enforce (ii), we use the transport plans *T*_*i*_ (1 *< i* ≤ *M*) as weighted links between the non-anchor datasets and the anchor so that strongly matched cells “pull” each other in the shared space.

#### Algorithm 1

**scSAGA**

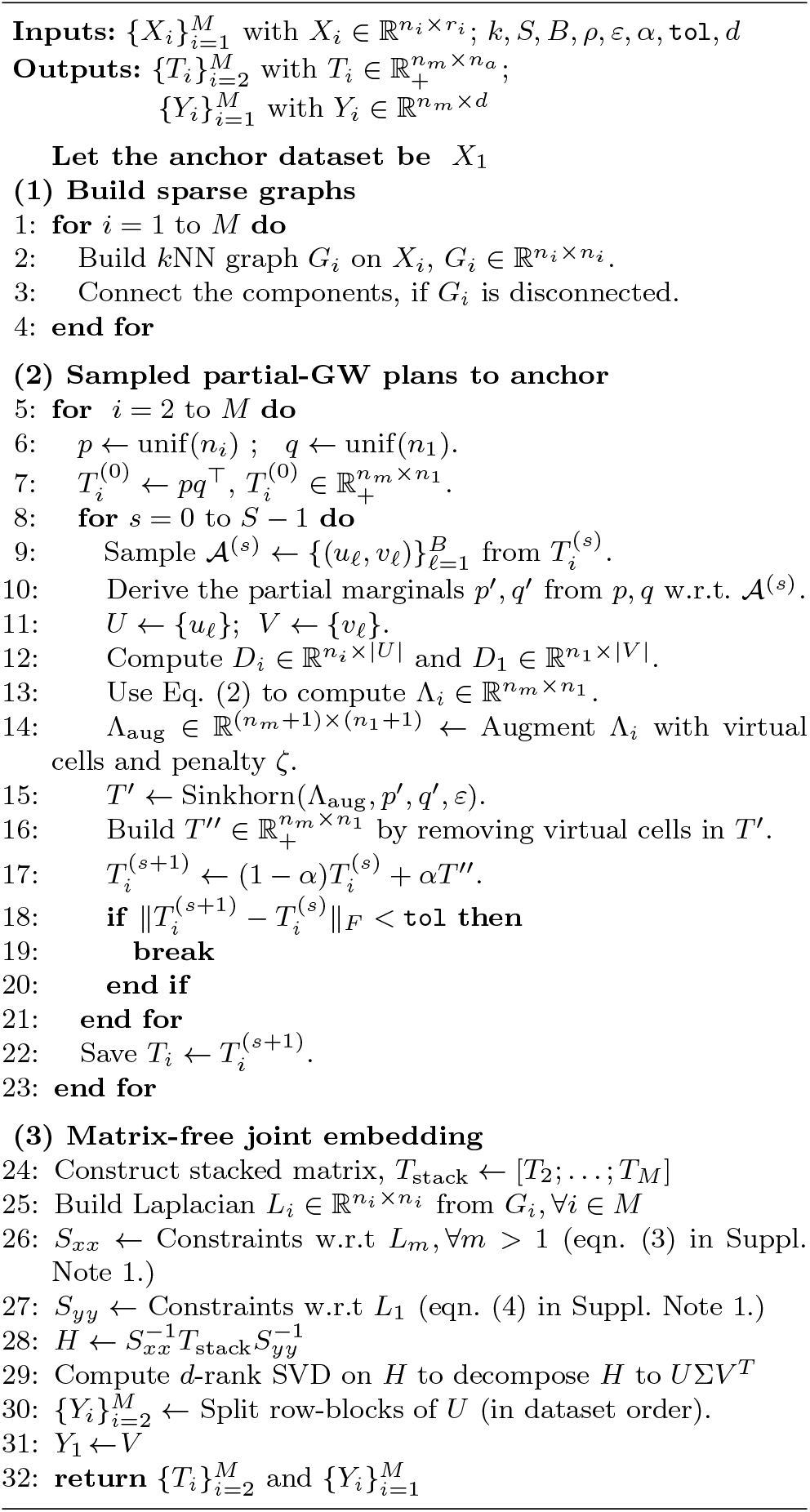

This approach is similar to prior GW-based pipelines that derive a shared space from transport plans (e.g., Pamona/SCOT), but the scSAGA approach implements them in a memory-efficient and matrix-free manner that can support integration of organism-wide datasets. Instead of constructing a large dense matrix for the full integrated system, we proceed the embedding computations as a sequence of sparse operations (graph-based smoothing plus transport-based averaging). Further, we use an iterative solver that only requires matrix–vector products (See Suppl. Note 1 for details).

### Implementation details

`scSAGA` is implemented in Python. Core computations use `NumPy`/`SciPy` for dense and sparse linear algebra, and `scikit-learn` [19] is used for *k*NN graph construction and stitching via `KDTree`. Entropically regularized (partial) GW subproblems are solved with the Python Optimal Transport library (`POT`) [12]. The sampled-GW loop computes geodesic distances on sparse graphs using scipy’s sparse implementation of Dijkstra’s algorithm, parallelized with `joblib`. We use `PyTorch` to enable parallel CPU/GPU tensor operations during GW optimization.

### Performance Assessment of scSAGA

#### Datasets

We evaluated scSAGA on three paired human datasets spanning a range of sizes: SNARE-seq (2,148 cells) [8], Human PBMC (22k cells) [2], and a larger human PBMC dataset (138k cells) [4]. Paired datasets provide ground-truth cell correspondences, thereby enabling direct assessment of matching accuracy. To test generality beyond human data, we additionally benchmarked on paired mouse alzheimer’s brain [3], paired Arabidopsis root [16], and paired zebrafish [1] datasets. Finally, we included unpaired datasets to evaluate modality mixing and scalability: supplementary table 1 summarizes all datasets, including 20 healthy PBMC datasets used in large-scale experiments ranging from ∼8k to 290k cells per dataset, with the largest integration totaling just over 1 million cells.

#### Environment setup

All experiments were run on a high-performance compute cluster, with each node equipped with a Dual Intel Xeon Gold 6226 CPU with 24 cores. Experiments on smaller datasets (less than 50k cells) were run on a system with 64GB RAM, while larger datasets were run on a system with 300GB RAM. Runs exceeding 14 hours were marked as “Did Not Finish” (*DNF*), and runs terminated by memory limits were marked as “Out Of Memory” (*OOM*).

#### Evaluation metrics

We evaluated the integrated datasets for both (A) integration quality and (B) biological conservation.

To evaluate the quality of integrated data, we use *Accuracy* and *Alignment Score*. **Accuracy** measures the matching correctness when ground-truth correspondences are available. For each cell *i*, we predict its match as arg max_*j*_ *T*_*ij*_ (transport plan *T*) and report the fraction of correctly matched pairs. **Alignment Score** quantifies neighborhood mixing across modalities in the joint embedding [7, 5]. Let 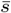be the average number of same-modality cells among each cell’s *k*NN. We compute Align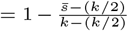, where 1 indicates perfect mixing and 0 indicates separation.

For measuring biological conservation, we used the following three metrics available with the SCIB software for evaluating single-cell integration [17]: **(i) Adjusted Rand Index (ARI)** measures agreement between clustering in the integrated space and cell-type labels (chance-corrected), with 1 indicating perfect agreement. **(ii) Normalized Mutual Information (NMI)** measures information overlap between predicted clusters and labels, ranging from 0 to

1. **(iii) Average Silhouette Width (ASW)** measures cluster compactness/separation. For cell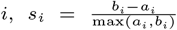, where *a*_*i*_ is the mean distance to same-type cells and *b*_*i*_ is the minimum mean distance to other types. We report 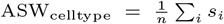, where higher values indicate better cell-type separation.

## Results and Discussion

### Quality assessment of scSAGA on paired and unpaired single cell datasets

We first benchmarked scSAGA on paired scRNA-seq/scATAC-seq human PBMC data with increasing integration sizes (600, 2k, 10k, and 22.6k cells), where known cell correspondences provide direct ground truth for evaluation. We compared scSAGA against two existing OT/GW-based methods – Pamona [7] and SCOT [10], and two other non-OT based methods - Seurat v5 [5, 21] and LIGER [25]. LIGER is also the only method available so far that can scale to large datasets.

Our results show that scSAGA achieves the best 1:1 matching accuracy across all human PBMC datasets (Table 1), remaining near-perfect on smaller size integrations and retaining high accuracy for larger size integrations. Amongst all the methods tested, SCOTv2 degrades most sharply as cell count increases, while the accuracy is lower for the other three methods by 18% - 32%, relative to scSAGA.

Two of the OT-based methods (scSAGA and Pamona) also achieve the strongest alignment scores (neighborhood mixing) on human PBMC across all dataset sizes (Table 2), while SCOTv2 and LIGER underperform and Seurat v5 trends downward for larger datasets. Taken together, our results show that scSAGA and Pamona are comparably strong methods in alignment, while scSAGA performs better in accuracy, and scales better (Tables 1–2).

**Table 2.**
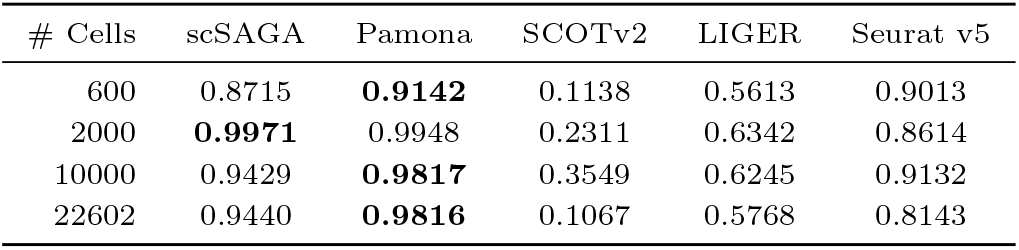
Alignment scores on the paired Human PBMC Granulocytes dataset for different integration sizes.

We next evaluated scSAGA’s integration quality on unpaired human PBMC scRNA-seq/scATAC-seq datasets with cells ranging from 17k to one million using the alignment score (Table 3). scSAGA maintains high alignment scores (0.75– 0.94) across all scales and completes every run. Pamona is competitive at 17k–37k but fails to complete runs on datasets beyond ∼37k. Similarly, SCOT also fails to complete runs for datasets with more than ∼37k cells. Seurat achieves reasonably good alignment scores for up to 450k cells (0.83–0.78), but fails to complete runs beyond that, while LIGER completes the full 1M-cell integration with consistently lower alignment scores (0.56–0.74) across the board. Notably, scSAGA successfully processes all datasets, achieving high alignment scores above 0.8 for all datasets up to 672k cells and a 0.75 score even at one million, indicating robust manifold preservation even at atlas scale.

**Table 3.** Alignment scores with unpaired multimodal single cell datasets. Comparison of scSAGA with baseline methods on progressively larger human PBMC integrations, from 17k up to 1M cells.

| Method | 17k | 37k | 162k | 252k | 414k | 450k | 672k | 1M | OOM Status |
| --- | --- | --- | --- | --- | --- | --- | --- | --- | --- |
| **scSAGA** | **0.9401** | 0.9257 | 0.8109 | **0.9107** | **0.8722** | **0.7819** | **0.8219** | **0.7523** | – |
| **Seurat** | 0.8710 | **0.9289** | **0.8324** | 0.7613 | 0.7839 | 0.7790 | – | – | Failed >450k |
| **Liger** | 0.7273 | 0.7410 | 0.6224 | 0.5611 | 0.6729 | 0.6916 | 0.6514 | 0.6877 | – |
| **Pamona** | 0.9312 | 0.8913 | – | – | – | – | – | – | OOM >37k |
| **SCOT** | 0.8115 | 0.7812 | – | – | – | – | – | – | OOM >37k |

scSAGA achieves superior accuracy and alignment by using kNN-graph geodesics to capture true manifold neighborhoods, whereas Pamona and SCOT often produce “blurry” alignments by spreading mass across multiple pairings. By employing plan-guided sampling and virtual-mass augmentation, scSAGA produces a sharper transport plan. Furthermore, its reliance on local graph structures rather than shared feature representations (like Seurat or Liger) ensures robust alignment in large, noisy, unpaired datasets.

### scSAGA is scalable and memory-efficient

To evaluate scalability, we benchmarked the runtime and peak memory utilized by scSAGA on paired and unpaired PBMC integrations ranging from hundreds of cells to one million, and compared against Pamona [7], SCOTv2 [10], Seurat v5 [21], and LIGER [25]. All runs were performed under identical conditions on 24-core CPUs with 64-300 GB RAM.

As shown in Fig. 2, scSAGA scales near-linearly in runtime (224 s at 22.6k; 2.98×10^3^ s at 160k; 2.4×10^4^ s at 1M) and in memory (0.14 GB at 600 cells up to 86 GB at 1M). This behavior is enabled by computing geodesic distances only for sampled anchors and using a matrix-free joint embedding implemented via sparse Laplacian solves. In contrast, Pamona and SCOT ran out of memory beyond ∼37k cells due to dense GW-style costs, while Seurat slowed substantially past ∼20k and failed beyond 450k. LIGER remains feasible at atlas scale but uses substantially more memory (e.g., 139 GB at 1M vs. 86 GB for scSAGA) and yields lower integration quality. The tabular version of these results is presented in Supplementary Table 2.

**Fig. 2.**
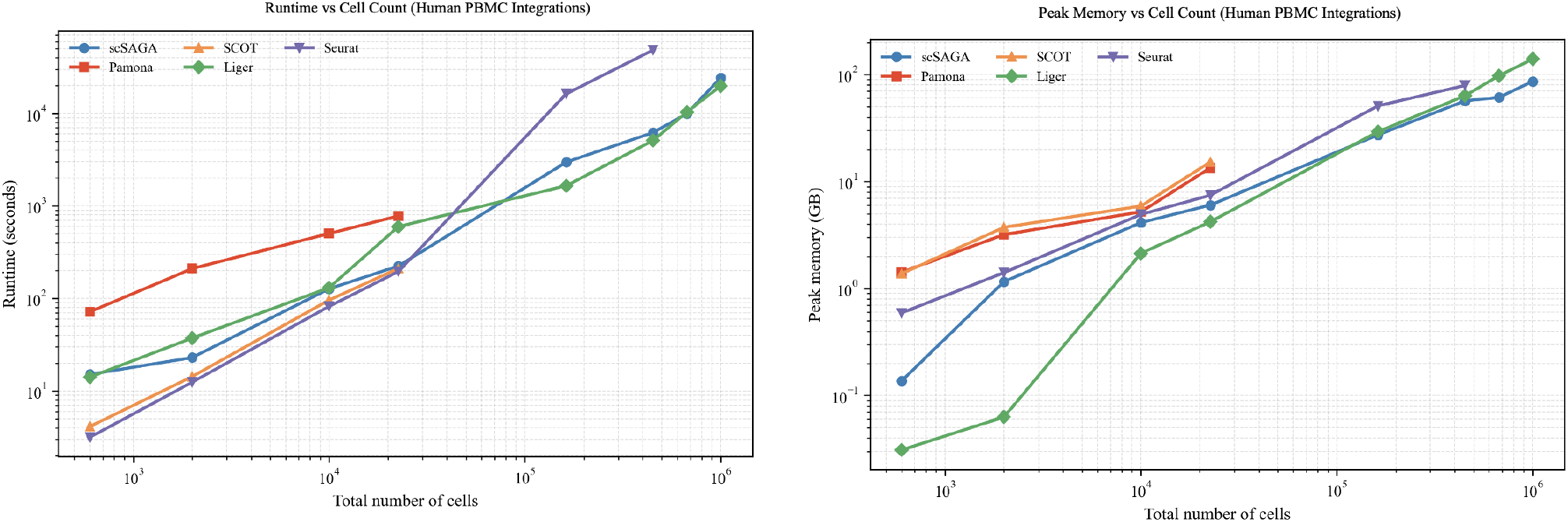
Runtime and memory consumption of various multimodal integration methods on single cell datasets with cells ranging from 17k to 1 million.

Together with the paired PBMC results (Tables 1–2), these benchmarks show that scSAGA simultaneously preserves integration quality and remains practical at large cell counts. This is mainly because our method avoids computing dense intra-domain distance matrices and computes the final joint embeddings sparsely, unlike Pamona and SCOTv2.

### Accurate single cell multimodal integration across various organisms with scSAGA

To assess generalization across divergent biological systems, we benchmarked scSAGA againt other methods on three multimodal datasets spanning distinct organisms: *Arabidopsis* root (48,948 cells) [16], mouse Alzheimer’s disease brain (66,918 cells) [3], human BMMC (138,498 cells) [4], and zebrafish neural cells (7871 cells) [1]. At full scale, Pamona and SCOT ran out-of-memory (OOM) on all three large size datasets, except for the Zebrafish dataset, consistent with the limited scalability of dense GW-style baselines. In contrast, scSAGA completed all integrations within 64 GB memory and achieved the strongest overall accuracy and alignment across organisms, relative to other methods (Tables 4–5).

**Table 4.** Cross-organism accuracy (1:1). “Full” uses the complete dataset; “20k” denotes subsampled datasets. OOM indicates out-of-memory. N/A indicates Not Applicable, as the dataset does not have 20k cells.

| Dataset | Method | Full | 20k |
| --- | --- | --- | --- |
| *Arabidopsis* root | scSAGA | **0.8600** | **0.9130** |
|  | Pamona | OOM | 0.8710 |
|  | SCOT | OOM | 0.6519 |
|  | LIGER | 0.4139 | 0.5302 |
|  | Seurat v5 | 0.0183 | 0.0500 |
| Mouse AD brain | scSAGA | **0.9100** | **0.9510** |
|  | Pamona | OOM | 0.8613 |
|  | SCOT | OOM | 0.8526 |
|  | LIGER | 0.7614 | 0.7577 |
|  | Seurat v5 | 0.7226 | 0.7934 |
| Human BMMC | scSAGA | **0.9310** | **0.9520** |
|  | Pamona | OOM | 0.8710 |
|  | SCOT | OOM | 0.8652 |
|  | LIGER | 0.8700 | 0.7900 |
|  | Seurat v5 | 0.8500 | 0.8342 |
| Zebrafish | scSAGA | **0.9714** | N/A |
|  | Pamona | 0.9632 | N/A |
|  | SCOT | 0.8950 | N/A |
|  | LIGER | 0.7398 | N/A |
|  | Seurat v5 | 0.0056 | N/A |

**Table 5.** Cross-organism alignment. “Full” uses the complete dataset; “20k” denotes subsampled datasets. OOM indicates out-of-memory. N/A indicates Not Applicable, as the dataset does not have 20k cells.

| Dataset | Method | Full | 20k |
| --- | --- | --- | --- |
| *Arabidopsis* root | scSAGA | **0.7989** | **0.9002** |
|  | Pamona | OOM | 0.8510 |
|  | SCOT | OOM | 0.5904 |
|  | LIGER | 0.3975 | 0.6100 |
|  | Seurat v5 | 0.0321 | 0.0100 |
| Mouse AD brain | scSAGA | **0.8931** | **0.9440** |
|  | Pamona | OOM | 0.8418 |
|  | SCOT | OOM | 0.8323 |
|  | LIGER | 0.5663 | 0.6953 |
|  | Seurat v5 | 0.7513 | 0.8430 |
| Human BMMC | scSAGA | **0.9200** | **0.9516** |
|  | Pamona | OOM | 0.8582 |
|  | SCOT | OOM | 0.8381 |
|  | LIGER | 0.8213 | 0.8321 |
|  | Seurat v5 | 0.8319 | 0.7915 |
| Zebrafish | scSAGA | 0.9320 | N/A |
|  | Pamona | **0.9463** | N/A |
|  | SCOT | 0.8880 | N/A |
|  | LIGER | 0.6534 | N/A |
|  | Seurat v5 | 0.0082 | N/A |

To enable a direct, matched comparison where all OT baselines complete runs, we additionally subsampled the large size datasets - *Arabidopsis* root, mouse Alzheimer’s disease brain, and human BMMC datasets - to 20,000 cells total using GeoSketch [13]. Under this fixed-size setting, all methods successfully ran, but scSAGA remained best overall across the three organisms, confirming that its advantage is not restricted to large-scale datasets (Tables 4–5 Notably, the largest performance gains are observed on the *Arabidopsis* and zebrafish datasets, where scSAGA substantially improves both accuracy and alignment relative to non-OT baselines, suggesting greater robustness to organism-specific feature differences, especially in comparison to Seurat. scSAGA does not assume a shared feature basis or rely on organism-specific feature engineering, which can degrade performance when regulatory programs and peak-to-gene relationships shift across species. Partial-GW further handles partial cell-type overlap by allowing unmatched mass, and kNN-geodesic costs keep geometry comparable even when raw feature spaces differ. As a result, Pamona remains competitive at smaller datasets because it is also geometry-based and utilizes partial-GW, while Seurat/LIGER can fail when shared-representation assumptions break, and SCOT/SCOTv2 can become diffuse due to full entropic GW forcing matches under incomplete overlap.

### scSAGA enables improved downstream clustering and annotation of single cell multimodal data

To further evaluate scSAGA’s utility for downstream biological interpretability, we employed 3 different evaluation measures from the scIB package [17] and quantified integration quality using Adjusted Rand Index (ARI), Normalized Mutual Information (NMI), and Average Silhouette Width (ASW_celltype_), computed between the integrated embeddings and known cell-type annotations. We compared scSAGA against Pamona, SCOT, LIGER, and Seurat using two different datasets: the human PBMC No Granulocyte dataset [2] and the SNARE-seq dataset [8].

Across both datasets (Figure 3), scSAGA consistently achieved the highest ARI, NMI, and ASW values, indicating accurate preservation of cell-type structure and improved cluster separability of cell-types in the integrated space. On the human PBMC dataset, scSAGA attained an ARI of 0.94, surpassing Pamona (0.85) and Seurat (0.89), while maintaining an ASW_celltype_ of 0.87. Additionally, on the SNARE-seq dataset, scSAGA achieved near-perfect scores (ARI = 0.96, NMI = 0.93, ASW = 0.92), reflecting strong alignment and biological coherence. In contrast, SCOT and LIGER exhibited reduced ARI and ASW, suggesting less accurate or over-smoothed integration. Overall, these results demonstrate that scSAGA yields embeddings that not only align modalities but also preserve biologically meaningful structure, leading to more reliable downstream clustering and cell-type annotation. Tabular values from figure 3 are given in suppl. table 3.

**Fig. 3.**
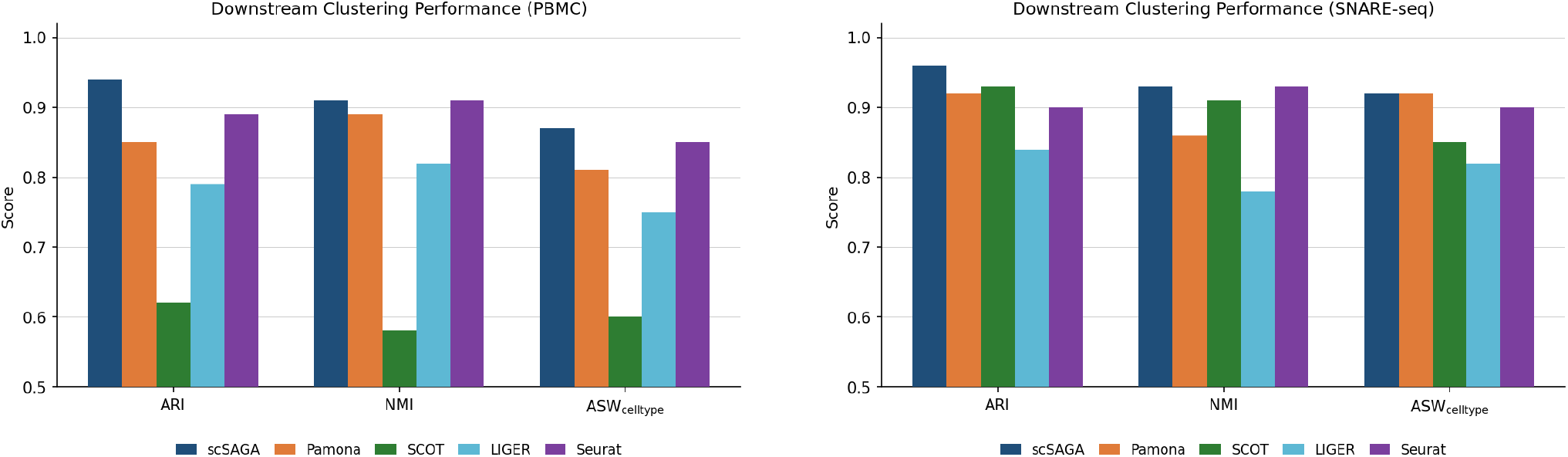
Downstream clustering and annotation performance. Higher ARI, NMI, and ASW indicate better agreement with true cell types and more coherent clusters. Results shown for (left) PBMC and (right) SNARE-seq datasets.

## Conclusions

We developed scSAGA, a scalable Gromov–Wasserstein (GW) alignment framework that makes geometry-preserving optimal transport practical for modern multimodal single-cell atlases. The central idea is to retain the geometric benefits of GW while removing its dense distance bottleneck: scSAGA represents each modality with a sparse neighborhood graph, evaluates GW using plan-guided sampling rather than all pairwise comparisons, and computes a shared embedding with a matrix-free solver that relies only on sparse operations.

Across paired and unpaired benchmarks spanning human PBMC/BMMC, mouse Alzheimer’s brain, Zebrafish and Arabidopsis root, scSAGA consistently achieved strong cross-modality mixing and higher matching accuracy than GW-based baselines (Pamona, SCOT) while remaining feasible at atlas scale where dense OT approaches fail. The resulting integrated spaces also improved the downstream clustering and annotation as measured by the SCIB metrics, indicating that scalability gains do not come at the expense of biological structure.

## Data Availability

Data used for all experiments in this paper are sourced from 10x Genomics and NCBI. The links to the raw data and all the accession numbers are listed in the supplementary table 1.

