## Supplemental Note 1 for "scSAGA: Single-cell Sampled Gromov Wasserstein Alignment for Scalable and Memory-efficient Integration of Multi-modal Single Cell Data"

### Supplemental Information

#### Supplementary Note 1: Matrix-free joint embedding in scSAGA

In this note, we discuss the complete detail Phase (3) of the scSAGA algorithm (lines **??** – **??**). In this phase, given the transport plans $T^{(i,1)}$ from each non-anchor dataset $i \neq 1$ to the anchor, we compute a shared low-dimensional embedding for all cells. This phase follows the same idea as Pamona: (i) Graph Laplacians are use to preserve local neighborhoods inside each dataset, and (ii) Transport plans enable connecting matched cells across datasets. The key contribution by scSAGA is that we compute in a matrix-free way, scaling to very large datasets. The phase proceeds as below:

1. Build neighborhood structure with graph Laplacians :
   For each dataset $i$, we construct graph Laplacian $L_i$ of the $k$NN graph $G^{(i)}$. The Laplacian encodes local neighborhoods: if two cells are neighbors in the original dataset, the embedding will encourage them to remain close. Laplacian can, thus be used, to preserve within-dataset local structure and geometry.

2. Convert transport plans into per-cell weights:
   Each transport plan $T^{(i,1)} \in \mathbb{R}^{n_i \times n_1}$ provides weighted links between cells in dataset $i$ and cells in the anchor dataset 1. We summarize the strength of participation of each cell in these links by taking simple sums of $T^{(i,1)}$.
   For each non-anchor dataset $i \neq 1$, we compute the row-sum vector $T^{(i,1)}\mathbf{1}$ and define
   $$\Sigma_x^{(i)} = \mathrm{diag}(T^{(i,1)}\mathbf{1}) \in \mathbb{R}^{n_i \times n_i}.$$
   Here, the $\ell$th entry of $T^{(i,1)}\mathbf{1}$ is the total transport mass leaving cell $\ell$ in dataset $i$ (its overall matching strength to the anchor). Placing these values on the diagonal gives a per-cell weight matrix for dataset $i$.
   For the anchor dataset, we compute column-sums and aggregate across all non-anchor datasets:
   $$\Sigma_y = \sum_{i \neq 1} \mathrm{diag}((T^{(i,1)})^\top \mathbf{1}) \in \mathbb{R}^{n_1 \times n_1}.$$
   The $j$th diagonal entry measures how much total transport mass is received by anchor cell $j$ from all other datasets. Thus, cells that are strongly involved in cross-dataset matchings receive larger diagonal weight.

3. Combine neighborhood preservation and transport weights :
   We then combine the Laplacians (neighborhood information) and the diagonal transport weights by defining
   $$S_{xx} = \mathrm{blkdiag}(L_i + \lambda \Sigma_x^{(i)})_{i \neq 1} \in \mathbb{R}^{N_{-1} \times N_{-1}} \quad (1)$$
   $$S_{yy} = L_1 + \lambda \Sigma_y \in \mathbb{R}^{n_1 \times n_1} \quad (2)$$
   where $N_{-1} \leftarrow \sum_{i \neq 1} n_i$, $\mathrm{blkdiag}(\cdot)$ stacks the non-anchor blocks along the diagonal and $L_1$ is the Laplacian of the anchor dataset. In words: the $L$ terms preserve local neighborhoods within each dataset, while the $\Sigma$ terms emphasize cells that are strongly supported by the transport plans. This is the same construction used in Pamona, adapted to the anchor setting.

4. Matrix-free use of $H$:
   We next define
   $$H = S_{xx}^{-1}\, T_{\text{stack}}\, S_{yy}^{-1}.$$
   This is similar to how Pamona performs the embedding pipeline: the transport plans $T_{\text{stack}}$ connect datasets, while $S_{xx}$ and $S_{yy}$ enforce neighborhood preservation and balance contributions using the transport mass.

Importantly, scSAGA *never explicitly forms* the matrix $H$ (or any large dense product). Instead, the rank-$d$ SVD is computed using an iterative routine that only needs products of the form $Hv$ and $H^\top v$ for vectors $v$. These products are computed by applying the factors in sequence: multiplying by $T_{\text{stack}}$, and solving linear systems with $S_{xx}$ and $S_{yy}$. Because each $L_i$ is sparse, each $\Sigma$ is diagonal, and $S_{xx}$ is block-diagonal, these operations can be done efficiently without constructing large dense matrices, which is why the embedding is matrix-free.

Finally, the left and right singular vectors of $H$ are returned as the shared embeddings.
